# Macrophages inhibit *Coxiella burnetii* by the ACOD1-itaconate pathway for containment of Q fever

**DOI:** 10.1101/2022.05.10.491306

**Authors:** Lisa Kohl, Md. Nur A Alam Siddique, Barbara Bodendorfer, Raffaela Berger, Annica Preikschat, Christoph Daniel, Martha Ölke, Michael Mauermeir, Kai-Ting Yang, Inaya Hayek, Manuela Szperlinski, Jan Schulze-Luehrmann, Ulrike Schleicher, Aline Bozec, Gerhard Krönke, Peter J. Murray, Stefan Wirtz, Masahiro Yamamoto, Valentin Schatz, Jonathan Jantsch, Peter Oefner, Daniel Degrandi, Klaus Pfeffer, Simon Rauber, Christian Bogdan, Katja Dettmer, Anja Lührmann, Roland Lang

## Abstract

Infection with the intracellular bacterium *Coxiella (C.) burnetii* can cause chronic Q fever with severe complications and limited treatment options. Here, we identify the enzyme cis-aconitate decarboxylase 1 (ACOD1 or IRG1) and its product itaconate as protective host immune pathway in Q fever. Infection of mice with *C. burnetii* induced expression of several anti-microbial candidate genes, including *Acod1*. In macrophages, *Acod1* was essential for restricting *C. burnetii* replication, while other antimicrobial pathways were dispensable. Intratracheal or intraperitoneal infection of *Acod1^-/-^* mice caused increased *C. burnetii* burden, significant weight loss and stronger inflammatory gene expression. Exogenously added itaconate restored pathogen control in *Acod1^-/-^* mouse macrophages and blocked replication in human macrophages. In axenic cultures, itaconate directly inhibited growth of *C. burnetii*. Finally, treatment of infected *Acod1^-/-^*mice with itaconate efficiently reduced the tissue pathogen load. Thus, ACOD1-derived itaconate is a key factor in the macrophage-mediated defense against *C. burnetii* and may be exploited for novel therapeutic approaches in chronic Q fever.

## Introduction

Q fever is an anthropozoonotic infection caused by *Coxiella (C.) burnetii*, mostly acquired by inhalation of infectious aerosol released from birth products of infected goats, sheep or cattle. Acute Q fever is characterized primarily by respiratory symptoms and pneumonia, but can also manifest as hepatitis (Eldin et al., 2017). In most patients, the disease is self-limiting and inflammation resolves within a few weeks. However, 1 to 5% of patients will develop chronic Q fever (Kampschreur et al., 2014), with persistent replication of *C. burnetii* in various tissues, including the vascular system. Severe complications such as endocarditis and mycotic aneurysms can lead to Q fever-related death in 25% of patients (van Roeden et al., 2019). Chronic Q fever is difficult to treat with antibiotics such as tetracyclines and fluoroquinolones, which show limited efficacy even after extended courses of 18 to 24 months.

*C. burnetii* is an intracellular bacterium that resists the harsh environment of the macrophage phagolysosome and in fact requires an acidic pH for replication. The bacterium expresses a type IV secretion system (T4SS) for injection of effector proteins into the host cell cytoplasm, which is required for formation of the *Coxiella-*containing vacuole (CCV) and intracellular replication (Carey et al., 2011; Luhrmann et al., 2017). Suppression of phagosome acidification by the drug hydroxychloroquine synergizes with the antibiotic doxycycline *in vitro* (Raoult et al., 1990). The combination of hydroxychloroquine with doxycycline is the currently recommended medical treatment for chronic Q fever (Anderson et al., 2013).

Studies in mouse models showed that immunologic control of *C. burnetii* requires CD4 and CD8 T lymphocytes and production of IFNγ (Andoh et al., 2007), the essential cytokine for arming macrophages to combat intracellular bacteria. Innate immune cells detect *C. burnetii* through Toll-like receptors (TLR) and MyD88-signaling (Ammerdorffer et al., 2015; Kohl et al., 2019; Ramstead et al., 2016; Zamboni et al., 2004). The effector mechanisms induced by IFNγ and MyD88 signaling to enable macrophages for the killing of *C. burnetii* are incompletely understood. In the past, a protective role has been demonstrated for the inducible or type 2 nitric oxide (NO) synthase (iNOS or NOS2) (Howe et al., 2002; Zamboni and Rabinovitch, 2003) that is strongly induced by combined TLR and IFNγ activation and generates NO from the amino acid L-arginine (Bogdan, 2015).

Among the large number of genes induced by IFNγ in myeloid cells, several other important antimicrobial proteins have been identified that mediate defense against various intracellular pathogens. These include 47 kDa and 65 kDa guanylate-binding proteins (GBPs) that are recruited to phagosomes containing *Mycobacterium (M.) tuberculosis* (Kim et al., 2011) and *Toxoplasma (T.) gondii* (Degrandi et al., 2013; Kravets et al., 2016; Yamamoto et al., 2012). Another example is the tryptophan-depleting enzyme indoleamine-2,3-deoxygenase (IDO) 1 which inhibits *Francisella (F.) tularensis* replication in the lung (Peng and Monack, 2010). While the function of GBP family members in infection with *C. burnetii* has not yet been addressed, a recent study reported that IDO1 impairs growth of *C. burnetii* in human THP1 macrophages (Ganesan and Roy, 2019).

During the past 10 years, a series of studies highlighted the importance of metabolic pathways in the host inflammatory response to infection (O’Neill and Pearce, 2016). Macrophages respond to TLR activation with a glycolytic switch (the so-called Warburg effect) and alterations in the tricarboxylic acid (TCA) cycle that include accumulation of succinate (Ryan and O’Neill, 2020). These changes in central energy metabolism of the cell also impact on the transcriptional regulation of gene expression in response to microbial danger, including enhanced expression of IL-1 (Ryan and O’Neill, 2020). The TCA cycle metabolite citrate plays an important role in immunity because it is essential for fatty acid synthesis and, in addition, can be metabolized to itaconate (ITA) that has immunoregulatory and anti-microbial functions (Williams and O’Neill, 2018). Citrate supports replication of *C. burnetii* and is an essential component of the axenic growth medium of *C. burnetii* (Omsland et al., 2009). In macrophages infected with *C. burnetii*, high intracellular citrate levels induced by the transcription factor STAT3 promote bacterial replication, whereas hypoxia inhibits STAT3 activation, citrate accumulation and *C. burnetii* growth through HIF1α (Hayek et al., 2019).

Itaconate (ITA) is derived from cis-aconitate, which is an intermediate in the conversion of citrate to isocitrate within the TCA cycle and serves as a substrate for the enzyme cis-aconitate decarboxylase (ACOD1) (encoded by *Acod1*, also termed immune responsive gene-1 [*Irg1*]). ACOD1 is associated with mitochondria (Degrandi et al., 2009) and catalyzes the decarboxylation of cis-aconitate to ITA (Michelucci et al., 2013). ITA has recently received a lot of attention for its immunoregulatory effects (Mills et al., 2018), but it can also inhibit the growth of several bacteria including *M. tuberculosis* (Michelucci et al., 2013) and *Legionella (L.) pneumophila* (Naujoks et al., 2016). More recently, the intracellular growth of *M. avium* (Gidon et al., 2021) and *Brucella species* (Lacey et al., 2021) was reported to be inhibited by the ACOD1-ITA axis.

In a recently established mouse model of infection with *C. burnetii* Nine Mile Phase II (NMII), we observed that efficient clearance of NMII from spleen, liver and lung was dependent on the TLR adapter protein MyD88 (Kohl et al., 2019). The increased bacterial burden seen in MyD88-deficient mice was accompanied by reduced expression of pro-inflammatory genes like CCL2 and IFNγ and less leukocyte infiltration into the tissues. Here, we identify the ACOD1-ITA axis as a key MyD88-dependent macrophage-autonomous determinant of early containment of *C. burnetii* and control of Q fever.

## Results

### ACOD1 belongs to a group of MyD88-dependent genes upregulated by C. burnetii infection in vivo

To dissect the mechanisms of MyD88-dependent killing of *C. burnetii*, we explored the expression of several genes with known or assumed anti-microbial function in the spleen of mice after intraperitoneal infection with NMII. Consistent with reduced IFNγ expression, the mRNA induction of *Nos2*, *Gbp1*, *Gbp2*, *Gbp4*, *Gbp5*, *Ido1* and *Acod1* was severely impaired in the spleens of *Myd88^-/-^* mice after intraperitoneal infection (Fig. 1A). Infection of primary murine and human macrophages *in vitro* rapidly triggered expression of *Acod1* (Fig. 1B, C). As *in vivo*, the MyD88 pathway was required for robust induction of *Acod1* in murine BMM, whereas TNF and type 1 IFN signaling, which both can enhance *Acod1* expression (Degrandi et al., 2009; Gidon et al., 2021; Shi et al., 2005), was not essential (Fig. 1D).

**Figure 1:**
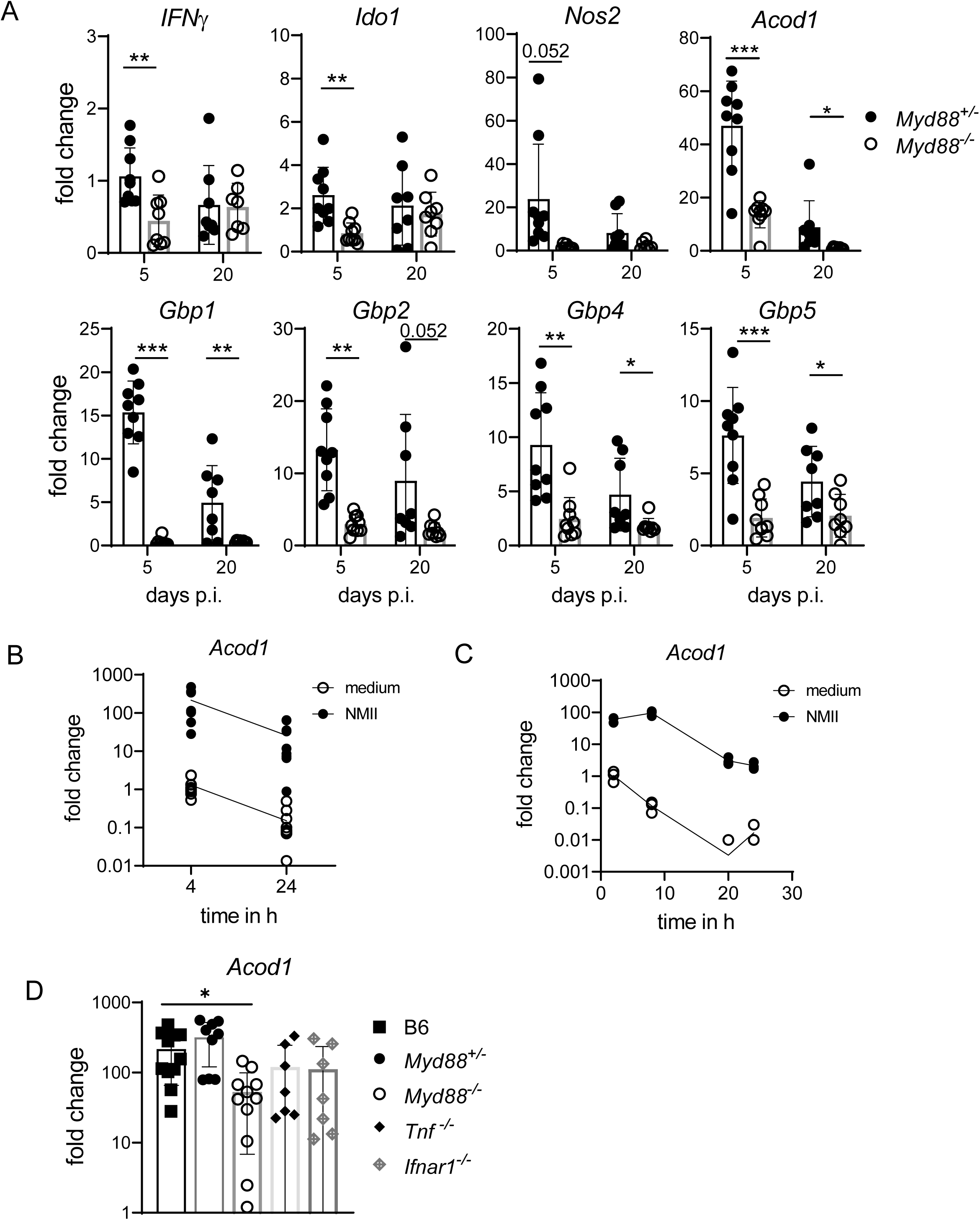
ACOD1 belongs to a group of MyD88-dependent genes upregulated by *C. burnetii* infection *in vivo* and *in vitro*. (A) *Myd88^+/-^* and *Myd88*^-/-^ mice were infected i.p. with NMII. Spleen RNA was prepared on day 5 post-infection. Expression of *Nos2*, *Gbp1*, *Gbp2*, *Gbp5*, *Ido1* and *Acod1* was analyzed by qRT-PCR, using non-infected *Myd88^+/-^* as calibrator. Mean and SD, n=8-9 mice per genotype, pooled from 3 experiments. Data for expression of *IFNγ*, *Nos2* and *Gbp1* on day 5 were previously reported (Kohl et al., 2019). (B) Induction of *Acod1* mRNA in mouse BMM infected with MOI 10 of NMII for 4 and 24 hours (fold change relative to mock controls, n= 9 mice pooled from 5 experiments). (C) *Acod1* mRNA expression in human GM-CSF-driven monocyte-derived macrophages (GM-MDM) infected with MOI 30 for 4 hours and harvested at the indicated times, n=3, one representative experiment of two performed. (D) *Acod1* expression in BMM is partially dependent on MyD88 but not on *Tnf* and *Ifnar1.* BMM were infected with NMII (MOI 10) for 4 hours. *Acod1* mRNA shown as fold induction relative to non-infected WT controls. Each dot represents one mouse, which were pooled from 3 to 5 independent experiments.

### ACOD1-deficient macrophages allow C. burnetii replication in vitro

We next employed bone marrow-derived macrophages (BMM) deficient in these candidate anti-microbial effector genes to test their requirement for control of NMII replication *in vitro* (Fig. 2A). A previous study reported that the production of NO by infected macrophages reduces the size of CCVs and thereby the intracellular load of *C. burnetii* (Zamboni and Rabinovitch, 2003). Using *Nos2^-/-^* macrophages, we observed only a weak, non-significant increase in the NMII burden as compared to wild-type controls (Fig. 2A). Several members of the guanylate-binding protein (GBP) family of IFNγ-inducible 65 kDa GTPases are recruited to the phagosomal/endosomal membrane in macrophages during infection with intracellular bacteria or protozoa and can cooperate to destroy intravacuolar pathogens. However, macrophages deficient in *Gbp2* or with a combined deletion of *Gbp1*, *Gbp2*, *Gbp3*, *Gbp5* and *Gbp7* encoded on chromosome 3 still efficiently inhibited NMII replication (Fig. 2A). IDO1 is the rate-limiting enzyme in tryptophan catabolism and has immunoregulatory as well as antimicrobial effects. Since tryptophan is an essential amino acid for *C. burnetii* (Sandoz et al., 2016), its intracellular depletion by IDO1 might constitute a nutritional control mechanism of macrophages as was recently shown in human macrophages (Ganesan and Roy, 2019). However, BMM harboring a combined deletion of *Ido1* and *Ido2* did not show any defect in restricting NMII replication (Fig. 2A). In contrast, macrophages from *Acod1^-/-^* mice were unable to restrict intracellular replication of NMII and harbored at least 10-fold more *Coxiella* after 120 hours (Fig. 2A), similar to *Myd88^-/-^* macrophages (Kohl et al., 2019).

**Figure 2:**
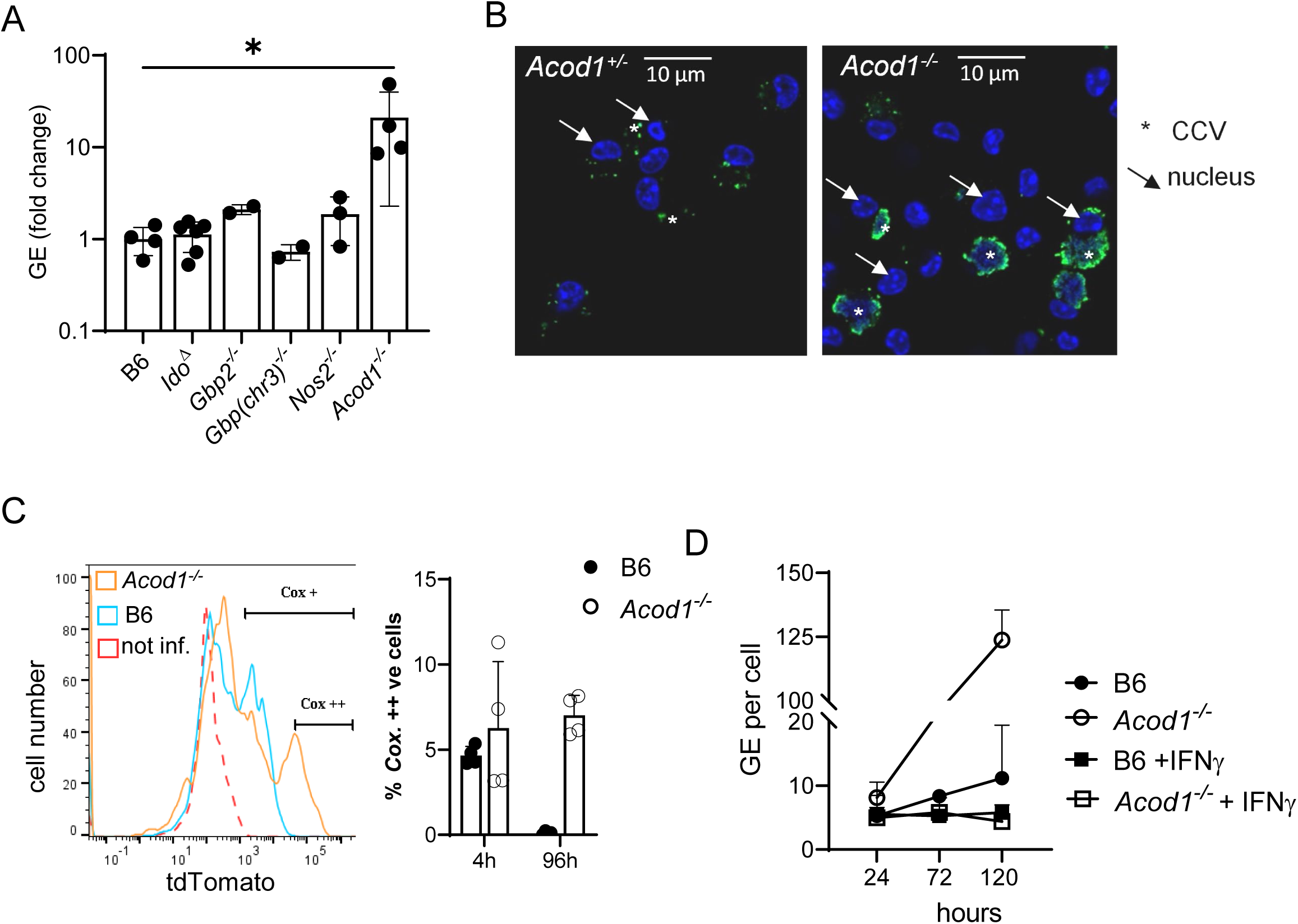
ACOD1-deficient macrophages allow *C. burnetii* replication *in vitro*. BMM of the indicated genotypes were infected with NMII (MOI 10) for 4 hours, followed by removal of extracellular bacteria by gentle washing. (A) Genome equivalents (GE) of NMII per cell were measured by qPCR after 120 hours in macrophage lysates. Average GE values from duplicate wells were normalized to the mean GE of all WT control samples. Each dot represents BMM from one mouse. * p<0.05 in Kruskal-Wallis test. (B) Immunofluorescence microscopy of WT and *Acod1^-/-^* BMM 120 hours after infection. Staining for *C. burnetii* appears in green, DAPI stain for DNA shows nuclei (examples marked by arrows) and large CCV in *Acod1^-/-^*BMM (marked by asterisk). (C) Flow cytometric detection of tdTomato- expressing NMII in BMM 4 and 96 hours after infection. Representative histogram overlay (after 96 hours) (*red* dotted line: WT, LPS; *blue*: WT, infected; *orange: Acod1^-/-^*, infected) and quantitation of highly tdTomato-positive BMM from 2 independent experiments. (D) Treatment with IFNγ overcomes the defect of ACOD1-deficient BMM to control NMII replication. Mean and SD of GE per cell of n=6 replicates (two mice per genotype, plated and infected in triplicate wells), representative of two experiments with similar results.

To validate the strong effect of ACOD1-deficiency on the abundance of *C. burnetii* genome equivalents detected by qPCR, we employed immunofluorescence staining. While wild-type macrophages showed only discrete staining with an anti-*C. burnetii* antiserum, large CCV were observed in *Acod1^-/-^* macrophages (Fig. 2B). Flow cytometry of macrophages infected with tdTomato-expressing NMII confirmed this effect in the absence of ACOD1, with around 6% of infected cells displaying a very strong fluorescent signal, corresponding to strongly enlarged CCVs filled with bacteria (Fig. 2C). While NMII strongly replicated in *Acod1^-/-^* BMM, treatment with IFNγ starting four hours after the infection, when extracellular NMII were washed away, inhibited bacterial replication in both WT and *Acod1^-/-^* BMM as shown by qPCR for *C. burnetii* GE (Fig. 2D) and immunofluorescence staining (Supplementary Figure S1). These data demonstrate that ACOD1 is essential for the cell-autonomous, but not for the IFNγ-driven control of *C. burnetii* growth.

### ACOD1-deficient mice have increased bacterial burden after infection with C. burnetii

We next asked whether ACOD1 was required for control of *C. burnetii* during infection *in vivo*. We infected mice by the intratracheal route with NMII. Seven days after infection, the bacterial burden in the lungs of *Acod1^-/-^* mice was more than 10-fold higher than in wild-type animals (Fig. 3A). Interestingly, this increased pulmonary bacterial replication was transient, as wild-type and *Acod1^-/-^*mice had comparable NMII numbers in the lung on day 11 (Fig. 3B). Immunohistochemistry for *C. burnetii* in the lungs confirmed a strongly increased bacterial load in *Acod1^-/-^* mice on day 7 after i.t. infection (Fig. 3C). As the NMII load in spleen and liver was not significantly different between the genotypes (Fig. 3A, B), we wondered whether ACOD1 was exclusively required for early control of *C. burnetii* in the lung in a tissue-specific manner, or whether the route of infection played a role. Infection of *Acod1^-/-^* and littermate *Acod1^+/-^*control mice by intraperitoneal injection revealed significantly higher bacterial loads in spleen, liver and lungs on day 7 (Fig. 3D), which was confirmed by immunohistochemistry of liver sections (Fig. 3E), demonstrating that ACOD1 is indeed required for the control of *C. burnetii* in all organs tested. Finally, to determine whether ACOD1 is necessary in hematopoietic or epithelial cells or in both compartments, bone marrow chimeric mice were generated and infected by intratracheal injection. The increased NMII burden in the lungs of irradiated WT mice reconstituted with *Acod1^-/-^* bone marrow, but not in the reverse setting, demonstrated a requirement for ACOD1 in hematopoietic cells (Fig. 3F). These findings, which are consistent with the reported predominant myeloid expression of *Acod1* (Wu et al., 2020), further underline a cell-autonomous effect of ACOD1 in infected macrophages.

**Figure 3:**
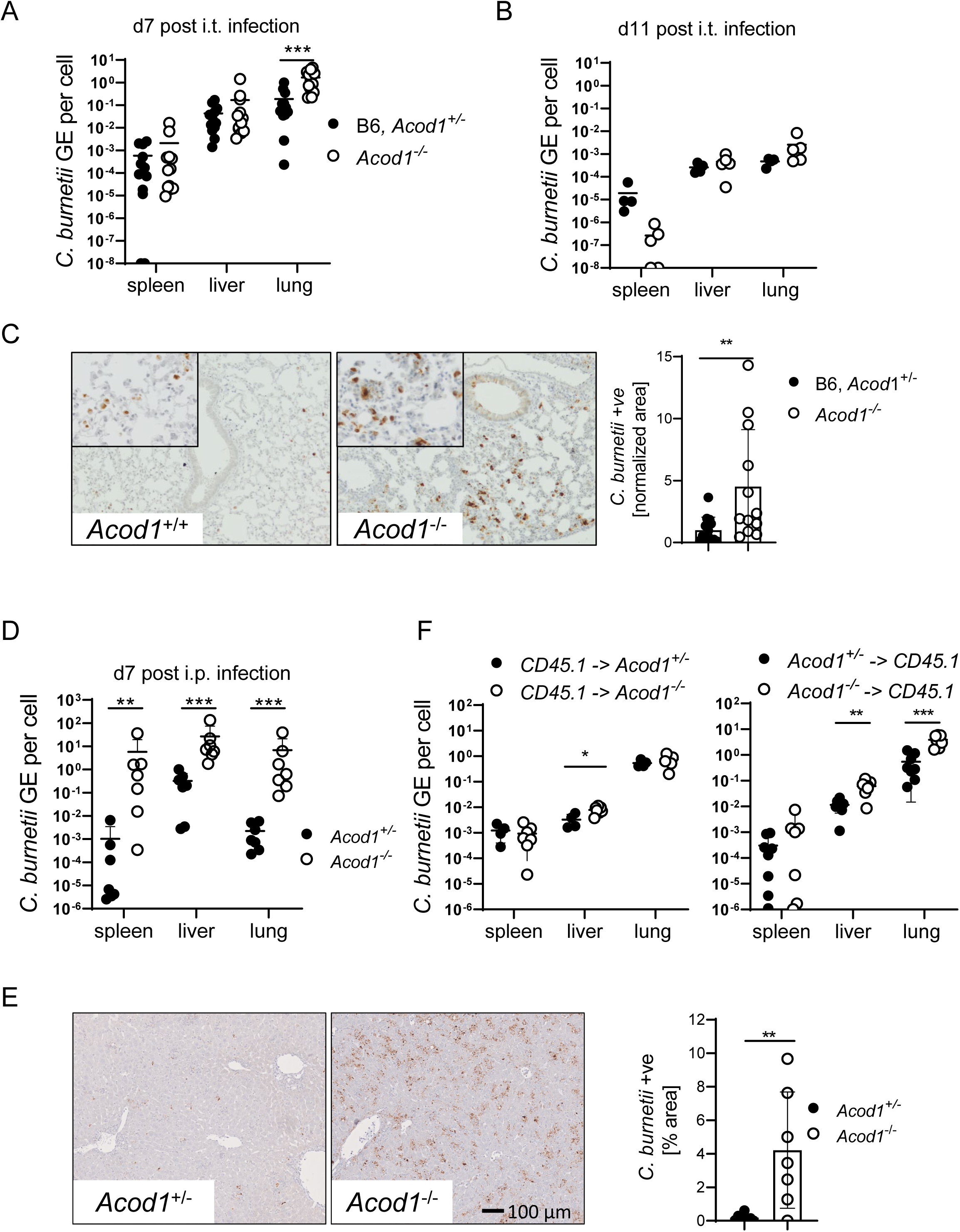
Increased bacterial burden in ACOD1-deficient mice infected with *C. burnetii*. Mice were infected by the i.t. route with 10^6^ NMII. Bacterial load in lung, spleen and liver on d7 (A) after i.t. infection. n=12 for *Acod1^-/-^,* n=13 for B6, *Acod1^+/-^* (pooled from three independent experiments). (B) Bacterial load on d11 after i.t. infection. n=5 for *Acod1^-/-^*, n=4 for B6. (C) Immunohistochemistry for *C. burnetii* in the lung on d7 after i.t. infection. Representative images and quantitation of lung tissue positive for *C. burnetii* (n=12-13 mice, pooled from three experiments, each normalized to the mean of WT or *Acod1^+/-^* infected mice). (D) Bacterial load in lung, spleen and liver on d7 after i.p. infection with 5x10^7^ NMII. t- tests adjusted for multiple testing using Holm-Sidak method, ** p<0.01, *** p<0.001. n=7 mice per genotype, pooled from two independent experiments. (E) Immunohistochemistry for *C. burnetii* on day 7 after intraperitoneal infection. (F) Radiation chimeric mice were infected i.t. with NMII. Each dot represents one mouse, data are pooled from two experiments. Bacterial burden in the tissues was determined after 7 days, except for one experiment with CD45.1 recipients when mice were sacrificed after 5 days because of weight loss. n=7-8 mice, * p<0.05 in Mann-Whitney test.

### Increased inflammatory and anti-microbial gene expression in ACOD1-deficient mice after C. burnetii infection

Intratracheal infection of *Acod1^-/^*^-^ mice led to a remarkable loss of body weight, which reached a maximum on day 7 after infection (Fig. 4A). Similar to the bacterial burden in the lung, the difference in weight was transient and disappeared until day 11. Following intraperitoneal infection, we also observed more pronounced weight loss in *Acod1^-/-^* mice (Fig. 4B). Deletion of *Acod1* in hematopoietic cells was sufficient for the development of an enhanced weight loss after i.t. infection with NMII (Fig. 4C).

**Figure 4:**
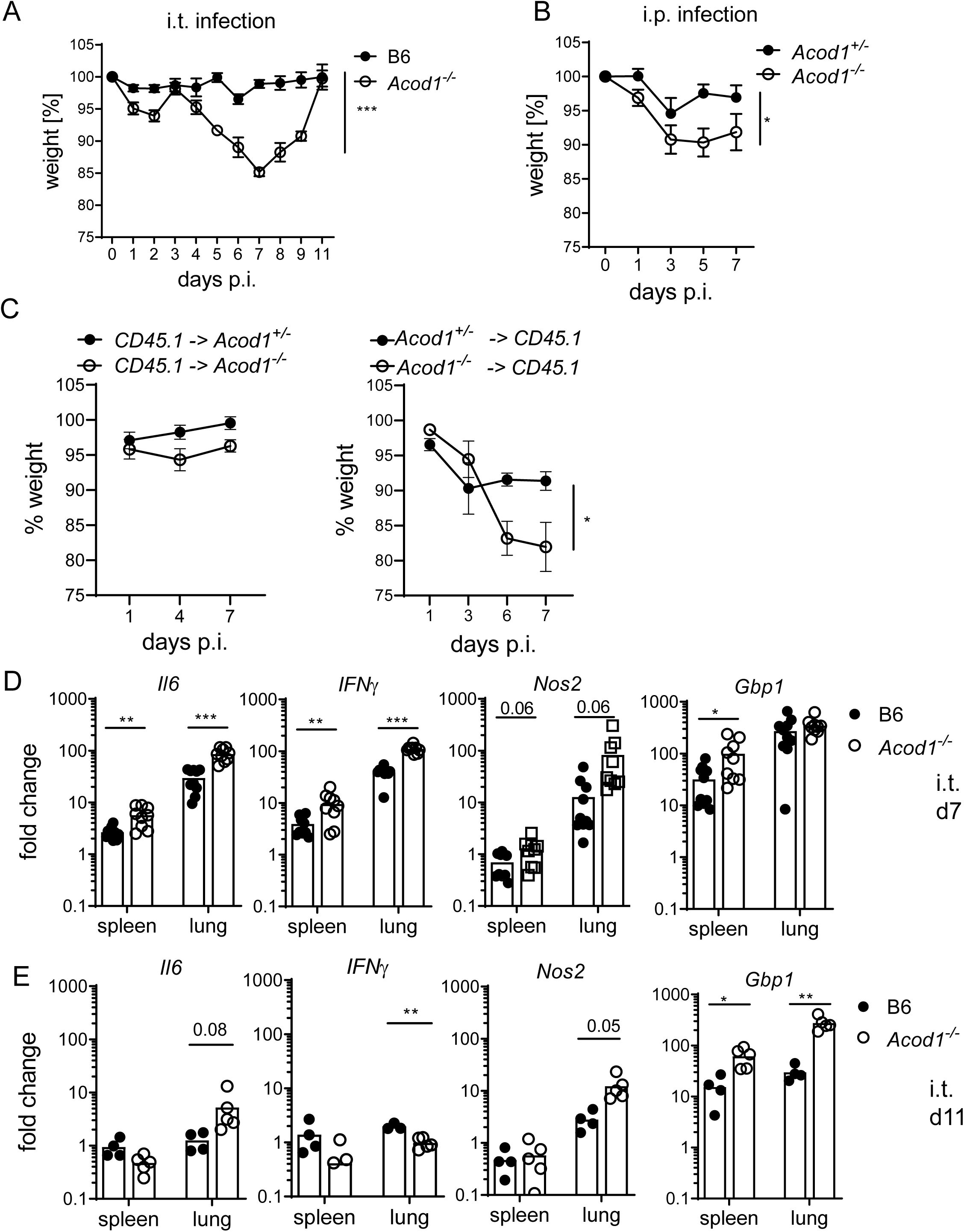
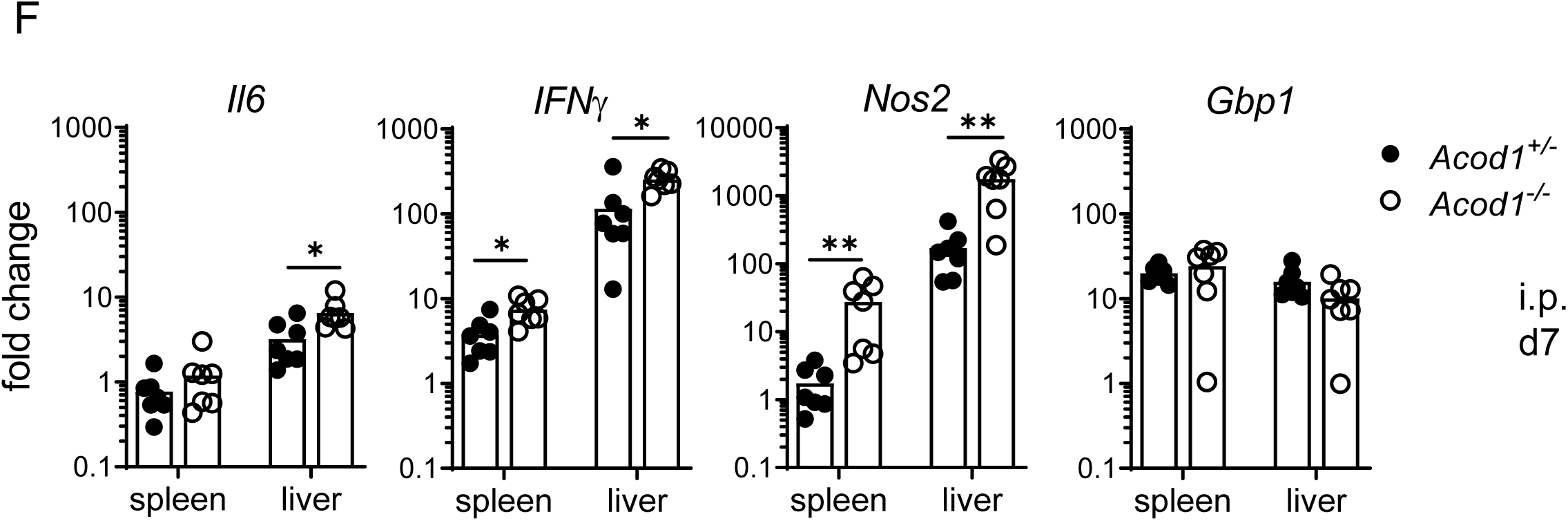
Weight loss and increased inflammatory and anti-microbial gene expression in ACOD1-deficient mice after *C. burnetii* infection. (A) Weight curve until d11 after i.t. infection. n=9-10 mice per group for days 1, 2, 4, 7; n=4-5 mice per group for days 3, 5, 8-11. (B) Weight curve after i.p. infection. Mean and SEM of n=7 mice per genotype. (C) Weight changes in the radiation chimeric mice. Mean and SEM of n=4-5 mice per group. Asterisks in (A-C) indicate * p<0.05, *** p<0.001 (2-Way ANOVA or mixed effects analysis). (D, E) Expression of *Il6*, *IFNγ*, *Nos2*, *Gbp1* in lung and spleen on d7 (D) and d11 (E) after i.t. infection is enhanced in the absence of ACOD1. Each dot represents one mouse. n=9-10, pooled from two experiments (D) and n=4-5 mice from one experiment (E). (F) Gene expression changes in spleen and liver after i.p. infection on d7. n=6-8 mice, pooled from two independent experiments. Asterisks in (D-F) indicate * p<0.05, ** p<0.01, *** p<0.001 (t-test comparing genotypes, adjusted by Holm-Sidak for multiple testing).

The phenotype of *Myd88^-/-^* mice was characterized by a high NMII burden and attenuated weight loss (Kohl et al., 2019) which illustrates that a higher *C. burnetii* load does not inevitably lead to an aggravation of clinical symptoms, which may instead correlate with the expression of inflammatory cytokines. Indeed, infected *Acod1^-/-^*mice expressed higher mRNA levels of *Il6*, *IFNγ*, *Nos2* and *Gbp1* than wild-type controls in the lungs, but also in the spleen, after intratracheal infection (Fig. 4D, E). A similar pattern of accentuated expression of *IFNγ* and *Nos2* was observed after intraperitoneal infection in spleen and liver, whereas *Gbp1* was not different between genotypes (Fig. 4F). Since NMII load was comparable in the spleen and livers of *Acod1^-/-^* and wild-type mice after intratracheal infection (Fig. 3A, B), these data indicate that ACOD1 exerts an immunoregulatory function as described previously in mice following endotoxin challenge or infection with *M. tuberculosis* (Mills et al., 2018; Nair et al., 2018). The enhanced production of IFNγ and of several of its anti-microbial target genes between day 7 and 11 after infection may explain the ability of *Acod1^-/-^* mice to resolve NMII infections despite an initial increase in bacterial replication.

### Exogenous itaconate restores intracellular ITA levels in ACOD1-deficient macrophages and restricts C. burnetii replication

To investigate the impact of *Acod1* deletion on macrophage ITA production and TCA cycle metabolites, we performed gas chromatography-mass spectrometry (GC-MS)-based analysis of mouse BMM infected with NMII (Fig. 5A). While ITA was hardly detectable (< 4 pmol/µg protein) in resting macrophages, NMII infection strongly increased the intracellular ITA levels (ca. 50 pmol/µg protein) in *Acod1^+/-^*but not in *Acod1^-/-^* macrophages. Consistent with the known inhibition of succinate dehydrogenase (SDH) by ITA (Cordes et al., 2016; Murphy and O’Neill, 2018), accumulation of succinate was observed in infected macrophages in an ACOD1-dependent manner. To test the assumption that increased replication of NMII in *Acod1^-/-^* macrophages is directly caused by a lack of ITA, we performed complementation experiments by adding exogenous ITA. Due to its polar nature, the capacity of ITA to pass the cell membrane and reach significant intracellular levels in macrophages has been questioned (Mills et al., 2018). We employed ITA itself and, in addition, its derivatives 4-octyl-itaconate (4-OI) and dimethyl-itaconate (DMI), which were previously used by others to achieve intracellular delivery (Mills et al., 2018; Zhang et al., 2020). Remarkably, ITA added at a concentration of 2.5 mM achieved high intracellular levels in both *Acod1^+/-^* and *Acod1^-/-^* macrophages, which in fact exceeded endogenous levels (Fig. 5A, upper panel). In contrast, neither addition of 4-OI (Fig. 5A) nor of DMI (data not shown) (both used at 250 µM) led to detectable ITA peaks, indicating that they were not converted to ITA intracellularly. These initially unexpected results on the uptake or intracellular delivery of ITA in macrophages are consistent with recent data from the Artyomov group that also did not find conversion of 4-OI or DMI to ITA in macrophages (Swain et al., 2020). Exogenous ITA, but not 4-OI or DMI, increased succinate also in *Acod1^-/-^* BMM (Fig. 5A). Intracellular levels of citrate were not altered by *Acod1* genotype or addition of ITA (Fig. 5A). Exogenous ITA efficiently enabled *Acod1^-/-^* macrophages to restrict replication of *C. burnetii* at concentrations of 1 and 2.5 mM (Fig. 5B), which showed no or little toxicity on macrophages based on MTT conversion assays (Supplementary Fig. S1). In contrast, 4-OI and DMI were highly toxic or did not significantly reduce NMII GE in *Acod1^-/-^* macrophages, respectively (Fig. 5B, Supplementary Fig. S2). Treatment of *Acod1^-/-^* macrophages infected by the tdTomato-expressing NMII strain with ITA prevented the formation of highly fluorescent cells containing large CCVs (Fig. 5C). Together, GC-MS analysis and addition of exogenous ITA demonstrated that uncontrolled replication of NMII in the absence of ACOD1 was directly due to a lack of intracellular ITA and not caused by other possible consequences of ACOD1-deletion such as the accumulation of citrate.

**Figure 5:**
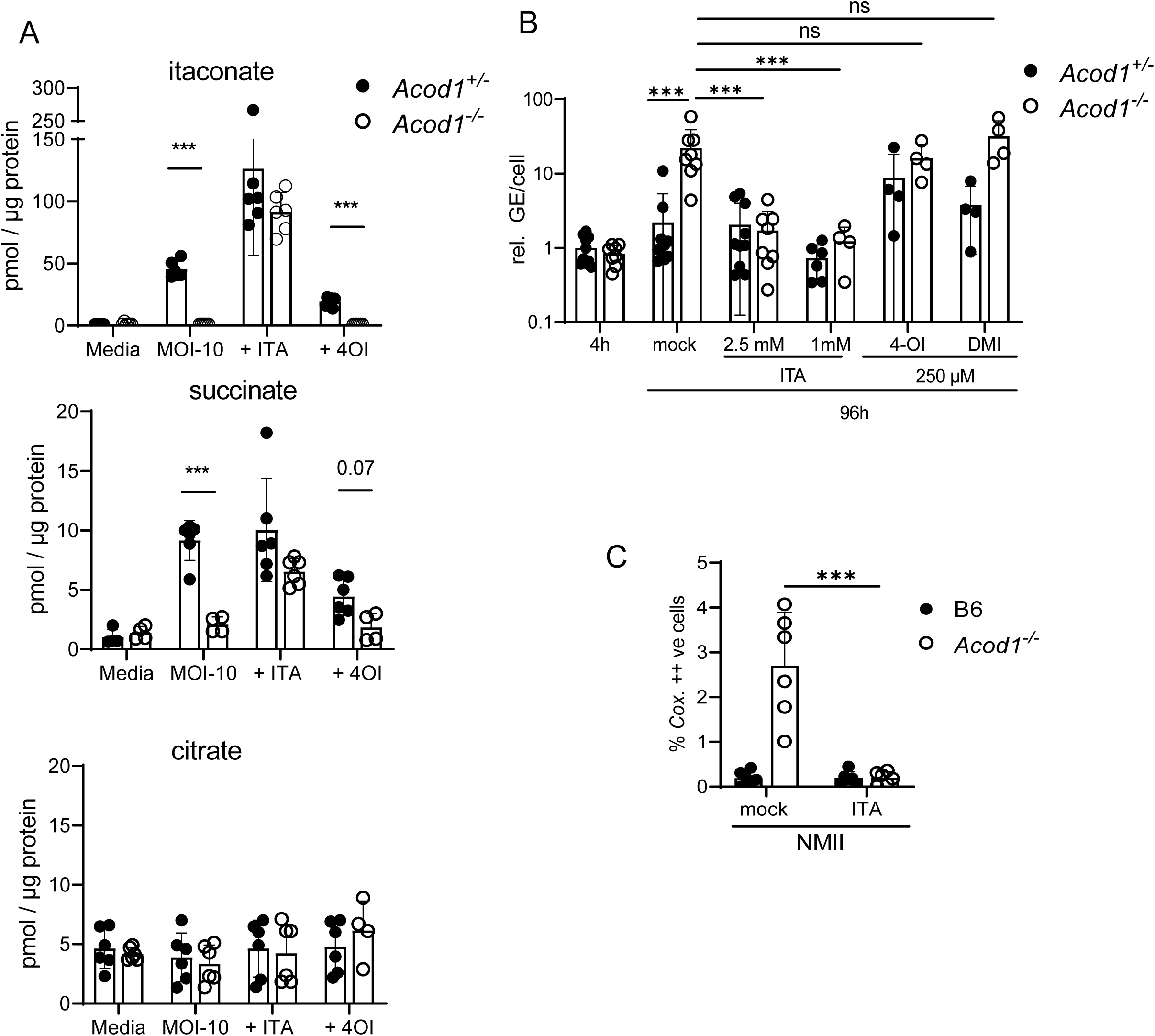
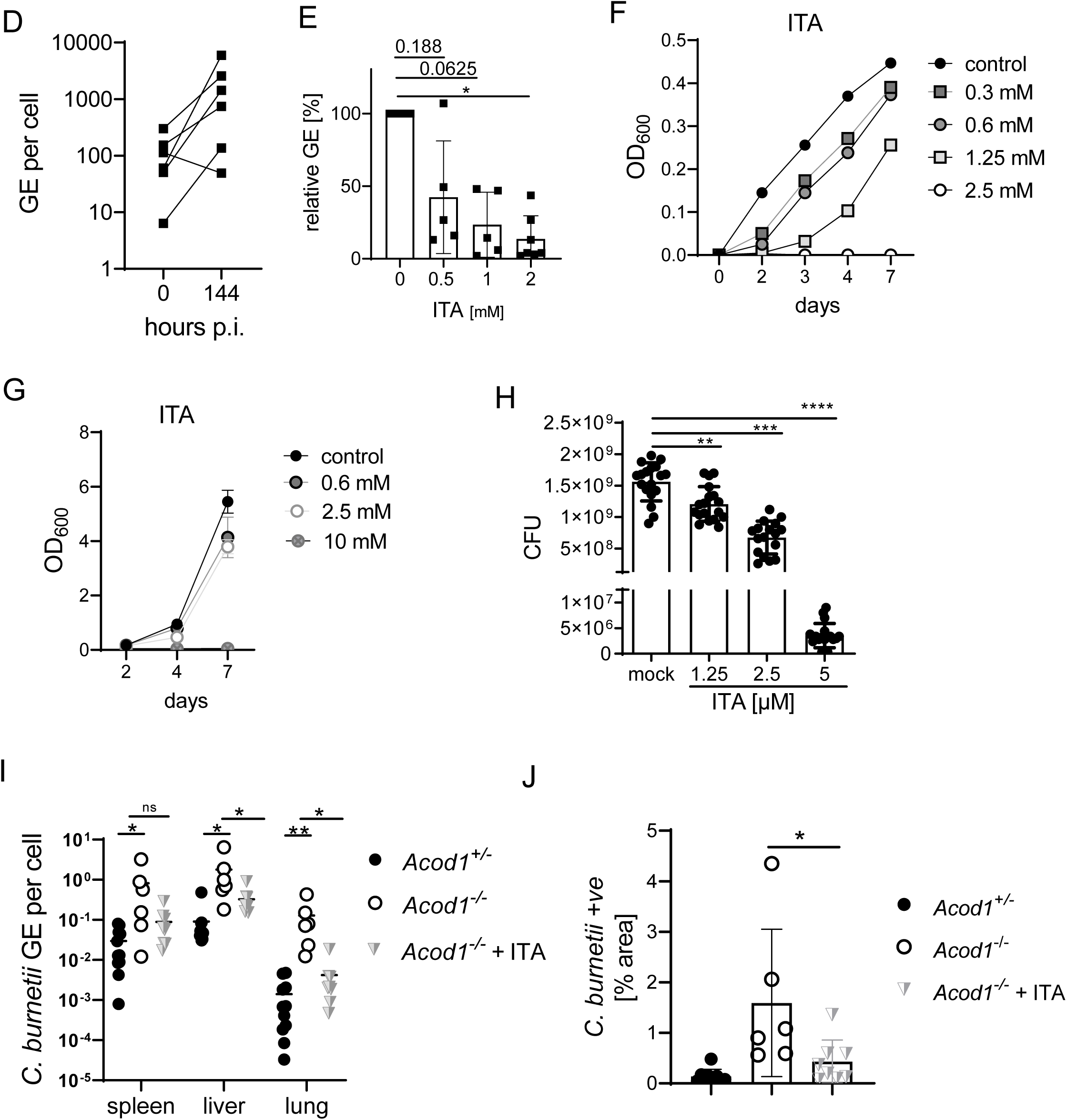
Exogenous ITA restores intracellular levels in ACOD1-deficient macrophages and restricts *C. burnetii* replication. (A) GC-MS analysis of TCA cycle metabolites and ITA in BMM infected with NMII (MOI 10). BMM were infected for 4h, followed by a washing step. Where indicated, exogenous ITA was added at 2.5 mM ITA, 4-OI at 250 µM, after removing extracellular bacteria. Macrophages were harvested after 24 h. Data shown are from two independent experiments, each performed in triplicate wells. (B) ITA, but not 4-OI or DMI, reduce NMII replication to WT levels in ACOD1-deficient BMM. Shown are qPCR data as *C. burnetii* GE normalized per cell (each dot represents one mouse and data were pooled together from 4 individual experiments with 2.5 mM ITA (n=8-9 mice per genotype) and two experiments with 1 mM ITA, 250 µM 4-OI and 250 µM DMI (n= 4 mice per genotype). Two-way ANOVA with Tukey’s multiple comparison test was performed in GraphPad Prism. (C) BMM were infected with tdTomato-expressing NMII and analyzed by flow cytometry 96 hour after infection. ITA was added at 2.5 mM. Depicted is the percentage of BMM highly positive for tdTomato (as in Fig. 2C). Pooled data from three independent experiments. (D) NMII replication in human GM-MDM. Infection was performed with MOI 10 for 4 hours followed by washing. Cells were lysed then directly or after 144 hours. Average values of replicate wells from 6 individual donors. (E) Addition of ITA inhibits NMII replication in human macrophages. ITA at the indicated concentrations was added to GM-MDM after infection with NMII MOI10. Bacterial burden was determined after 144 hours as GE per cell by qPCR. Data were normalized to GM-MDM infected but not treated with ITA (100 %). Average values of replicates from n=5-7 donors. p-values (Wilcoxon matched-pars signed rank test) are indicated. (F) ITA inhibits NMII replication in axenic culture in ACCM-2. Dose response and kinetic analysis. (G) Effect of ITA on *Mycobacterium bovis* BCG. Inoculum density on d0: OD_600_ = 0.045. Mean and SD, n=3. (H) ITA inhibits NMII growth in axenic culture after delayed addition of ITA at the indicated concentrations on day 5. On day 7, CFU were determined. Data points show results pooled from two independent experiments with three biological and technical replicates each. Mann-Whitney test comparing to mock. (I, J) Treatment of mice with 1 mg of ITA i.p. on days 1, 3, and 5 after i.p. infection with NMII lowers bacterial burden in the organs. *C. burnetii* GE quantified by qPCR (I), quantification of immunohistochemistry for *C. burnetii* in liver sections as in Fig. 3E (J). n=6-9 mice (for NMII infections), pooled from two experiments. Asterisks indicate significance (ANOVA with Dunnett’s multiple comparison in (I), Mann-Whitney test in (J).

We next asked whether the ACOD1-ITA pathway is also relevant for the control of NMII in human macrophages. A 10-fold increase of the number of intracellular bacteria was observed 144 hours after infection of human macrophages with NMII (Fig. 5D), which was blocked by the addition of 0.5 to 2 mM ITA (Fig. 5E). ITA may interfere with the replication of intracellular bacteria through effects on host cell metabolism, such as the inhibition of the mitochondrial complex II in the case of *F. tularensis* (Jessop et al., 2020); ITA can also act directly on bacteria, as demonstrated for *M. tuberculosis* by inhibition of the glyoxylate shunt enzyme isocitrate lyase (Michelucci et al., 2013). Therefore, we determined whether ITA impaired directly the replication of NMII. Addition of ITA to axenic cultures of NMII showed a dose-dependent inhibition of bacterial growth at concentrations between 0.3 and 2.5 mM (Fig. 5F), which are comparable to those found to inhibit replication in infected macrophages (Fig. 5B). NMII was more sensitive to inhibition by ITA than *M. bovis* BCG whose growth was inhibited only at concentrations above 2.5 mM (Fig. 5G). Of note, ITA did not only prevent replication of NMII when added at the beginning of the culture, but it also reduced bacterial numbers when used to treat established cultures between day 5 and day 7 (Fig. 5H). Importantly, while ITA at 1.25 and 2.5 mM inhibited further bacterial replication, the higher concentration of 5 mM ITA reduced the CFUs by a factor of more than 300, indicating bactericidal activity (Fig. 5H).

Given the strong direct activity of ITA against NMII, its effectiveness to inhibit intracellular replication in macrophages *in vitro*, and its low toxicity *in vivo* (Cordes et al., 2020), we finally asked whether treatment of mice with ITA can reduce *C. burnetii* burden *in vivo*. Treatment of *Acod1^-/-^* mice infected with NMII with 1 mg ITA injected i.p. every other day significantly reduced the bacterial burden in liver and lung as determined by qPCR for *C. burnetii* genome equivalents (Fig. 5I) or by immunohistochemistry (Fig. 5J).

## Discussion

This study shows that *C. burnetii* infection strongly induces *Acod1* expression which turned out to be essential for the control of NMII replication in macrophages *in vitro* and *in vivo*. Notably, the effect of deletion of *Acod1* on *C. burnetii* replication was much stronger than the impact of deficiencies of several other IFNγ-induced antimicrobial defense mechanisms (NOS2, IDO1/2, GBPs), thus establishing ACOD1 as a major protective determinant in the host defense against Q fever.

It is well known that ACOD1 catalyzes the generation of ITA by decarboxylation of cis-aconitate. Our results from GC-MS analyses showed that *Coxiella*-induced *Acod1* expression in macrophages led to high-level ITA production, which was absent in *Acod1*-deficient cells. Thus, ACOD1-mediated ITA generation appears to be required for curbing *C. burnetii* growth in macrophages. Since the first report of inhibition of *M. tuberculosis* by ITA (Michelucci et al., 2013), a similar effect has been shown for other intracellular bacteria, including *M. avium* (Gidon et al., 2021), *L. pneumophila* (Naujoks et al., 2016), *Salmonella (S.) typhimurium* (Chen et al., 2020), *Brucella melitensis* (Demars et al., 2021; Lacey et al., 2021) and *F. tularensis* (Jessop et al., 2020), but also for the extracellular bacteria *Escherichia (E.) coli* (Duncan et al., 2021) and *Staphylococcus (S.) aureus* (Singh et al., 2021). Remarkably, the ITA concentrations that we found to be required for inhibition of *C. burnetii* growth in macrophages or axenic cultures (0.6 – 2.5 mM), were lower than those documented for the above-mentioned pathogens (> 5 mM).

The mechanism of action for the antibacterial activity of ITA is incompletely understood and appears to be diverse.

Firstly, ITA can act indirectly by altering cellular metabolism of the host cell and promote antimicrobial effector functions, such as mitochondrial ROS production (Hall et al., 2013). In the case of *F. tularensis*, even high concentrations of ITA (50 mM) were ineffective in axenic culture, but inhibition of mitochondrial complex II (which is identical to SDH) by 4-OI and dimethyl-malonate in macrophages was associated with inhibition of *F. tularensis* replication (Jessop et al., 2020). We have confirmed that ACOD1-mediated ITA production leads to high intracellular succinate levels in macrophages infected with *C. burnetii*, but have not addressed whether ROS production was affected in our system.

Secondly, ITA can be directly inhibitory to bacterial replication through different mechanisms. The enzyme isocitrate lyase (ICL) is required for the glyoxylate pathway and is a target for ITA in *M. tuberculosis* (Michelucci et al., 2013) and *B. melitensis* (Demars et al., 2021). In addition, the ITA metabolite itaconyl-CoA inhibits methylmalonyl-CoA mutase of *M. tuberculosis* to interfere with propionate-dependent growth (Ruetz et al., 2019). For other bacteria, the mechanism of growth inhibition by ITA is unknown to date. Given the acidic conditions in the phagolysosome, the recent demonstration of synergy between ITA and low pH for blocking replication of *E. coli* and *S. typhimurium* (Duncan et al., 2021) is of interest.

Several lines of evidence in our results indicate that ITA generated in infected macrophages directly inhibits the replication of *C. burnetii*. Firstly, exogenous ITA provided to *Acod1^-/-^* macrophages at non-toxic concentrations complemented intracellular ITA levels and inhibited NMII growth within macrophages. Secondly, deficiency in *Acod1* and exogenous ITA did not alter the intracellular levels of citrate, making indirect effects through accumulation or depletion of this essential substrate for *C. burnetii* growth unlikely. Thirdly, ITA impaired the growth of NMII in axenic culture media at concentrations between 0.6 and 2.5 mM, which are much lower than for the ITA-controlled pathogens mentioned above and are easily achieved intracellularly in activated and bacteria-infected murine macrophages (Chen et al., 2020).

As ACOD1 is localized in mitochondria, newly generated ITA needs to reach the CCVs to exert its effect. In *Salmonella*-infected macrophages, this transport is facilitated by the GTPase Rab32 that interacts with ACOD1 and is required for delivery of ITA to the *Salmonella*-containing vacuole (Chen et al., 2020). It is likely that positioning of ACOD1 in the vicinity of the pathogen-containing vacuole results in even higher local concentrations so that a direct bactericidal effect is conceivable as observed here for *C. burnetii* in axenic culture with 5 mM ITA. *C. burnetii* does not utilize the glyoxylate shunt and lacks the enzyme ICL (Seshadri et al., 2003). Therefore, the exact molecular mechanism of the inhibitory effect of ITA on *Coxiella* remains to be elucidated.

The immunoregulatory activity of ACOD1-ITA in sepsis and inflammation models has received more attention than its antimicrobial effects (Mills et al., 2018; Nair et al., 2018; Swain et al., 2020). *Acod1^-/-^*mice infected with NMII showed significant weight loss and increased inflammatory gene expression compared to B6 or *Acod1^+/-^* controls. These differences are difficult to interpret, as they may be caused by the increased bacterial load. However, two observations indicate that ACOD1 not only shows direct antimicrobial activity on *C. burnetii* but indeed has immunoregulatory effects in Q fever: firstly, we previously found in *Myd88^-/-^* mice that increased bacterial burden does not *per se* lead to stronger inflammatory responses; and secondly, the NMII burden in the spleen after intratracheal infection was not increased in *Acod1^-/-^*mice, but expression of *Il6*, *IFNγ* and *Gbp1* was still significantly higher (Fig. 4D). Thus, although we interpret the unimpeded growth of NMII during the first week of infection as the major driver of inflammation and clinical disease in *Acod1^-/-^*mice, the loss of immunoregulatory functions of ACOD1 may contribute.

The increased *C. burnetii* burden and weight loss after intratracheal infection was transient and control of infection after day 7 coincided with the expected generation of adaptive immunity. The strong expression of IFNγ-induced antimicrobial programs, as indicated by the robust upregulation of *IFNγ*, *Nos2* and *Gbp1* in the mice on day 7 and day 11, likely compensates for the lack of ITA in *Acod1^-/-^* mice at this later phase of infection. This notion is supported by our observation that stimulation of *Acod1^-/-^*BMM with IFNγ enables them to control the growth of *C. burnetii* (Fig. 2D).

Treatment with ITA in mice was recently shown to improve the outcome in a reperfusion injury model (Cordes et al., 2020) and to counteract lung fibrosis when inhaled (Ogger et al., 2020). In addition, administration of DMI protected from brain injury in cerebral ischemia (Kuo et al., 2021). Further, application of 4-OI in a murine endophthalmitis model caused by *S. aureus* infection synergized with antibiotics to reduce bacterial burden and attenuated inflammatory damage (Singh et al., 2021). The successful treatment of *Acod1^-/-^* mice with intraperitoneal ITA injections shown here demonstrates that sufficiently high concentrations can be achieved *in vivo* to inhibit the growth of *C. burnetii*. While in wild-type mice, the high levels of endogenously produced ITA may not be further increased by ITA treatment, the situation is probably different in humans. Based on published results (Michelucci et al., 2013) and our own preliminary data (not shown), human macrophages produce much less ITA than their murine counterparts. This species-difference in ITA generation might explain why exogenous ITA was effective in reducing NMII burden in our experiments with human MDM (Fig. 5F), whereas no inhibition was observed in *Acod1*-expressing murine macrophages (Fig. 5B). If this low-level of ITA in humans compared to mice also occurs *in vivo* during infection, exogenous ITA would be expected to boost the capacity of infected macrophages to control *C. burnetii*. Thus, our results suggest that ITA should be further explored as a therapeutic strategy in acute and chronic Q fever.

## Materials and Methods

### Reagents

Itaconate (ITA) and its derivatives 4-octyl itaconate (4-OI) and Dimethyl itaconate (DMI) were purchased from Sigma-Aldrich (Deisenhofen, Germany) (ITA, I29204-100G; 4-OI, SML2338- 25mg; DMI, 592498-25G) as powder. Stock solutions were prepared in PBS (ITA and DMI) or DMSO (4-OI).

### Mice

All mice utilized in this study were at least six weeks of age. C57BL/6 wild-type mice were obtained from Charles River Breeding Laboratories (Sulzfeld, Germany) or bred at the Präklinische Experimentelle Tierzentrum of the University Hospital Erlangen (PETZ). *Acod1^-/-^* and *Nos2^-/-^* mice were from Jackson Laboratory (C57BL/6NJ-*Acod1^em1(IMPC)J^/*J and B6.129P2- *Nos2^tm1Lau^*/J)). *Tnf^-/-^* mice (Korner et al., 1997) were obtained from Dr. H. Körner (University of Tasmania, Australia), *Ifnar1^-/-^* mice (Muller et al., 1994) from Dr. U. Kalinke (TWINCORE, Hannover, Germany). *Myd88^-/-^* mice (*Myd88^tm1Ak^*^i^) were provided by Dr. S. Akira (University of Osaka, Japan) (29). *Gbp2^-/-^* mice (*Gbp2^tm1.1Kpf^*) were provided by Dr. K. Pfeffer (Universitätsklinikum Düsseldorf, Germany) (Degrandi et al., 2013). Gbp(chr3) mice lacking *Gbp1*, *Gbp2*, *Gbp3*, *Gbp5* and *Gbp7* (Del(3Gbp2-ps-Gbp5)1Ktak) (Yamamoto et al., 2012) were obtained from Dr. Thomas Henry (University of Lyon, France). Mice with a combined deletion in *Ido1* and *Ido2* (*Ido1-Ido2^ΔΔ^*) (Van de Velde et al., 2016) were generated and provided by one of us (P.M.). All mice were generated on a C57BL/6 background or crossed back multiple generations to C57BL/6.

### *Culture of* Coxiella burnetii

An isolate of the *C. burnetii* Nine Mile phase II (NMII) strain clone 4 (NMII, RSA493) was generously provided by Matteo Bonazzi (Institut de Recherche en Infectiologie de Montpellier, Montpellier, France). One aliquot of purified NMII was propagated in a 75 cm^2^ tissue culture flask containing 30 mL acidified citrate cysteine medium (ACCM-2, 4700-003, Sunrise Science Products, San Diego, CA, USA) at 37°C in an humidified atmosphere of 5% CO_2_ and 2.5% O_2_ (Omsland et al., 2009). After four days of culture, NMII were transferred overnight to room temperature and ambient atmosphere. Subsequently, bacteria were pelleted for 15 min at 4,500 x g, resuspended in 1 mL phosphate buffered saline (PBS) and quantified by optical density at OD_600_, where an OD_600_ of 1 equals 1 x 10^9^ NMII per mL. Bacteria were diluted in PBS or cell culture media without antibiotics and kept on ice until use for infection of mice or macrophages.

### Generation of tdTomato-expressing C. burnetii

The tandem-di-Tomato (tdTomato) expressing NMII strain was prepared as described previously (Matthiesen et al., 2020): the tdTomato gene (codons optimized for expression in *C. burnetii*) was amplified by PCR with Q5 polymerase (New England Biolabs, Frankfurt, Germany) using primers a533 (5’-GATTTAAGAAGGAGATCTGCAGATGGTGTCAAAAGGAG-3’) and a534 (5’-AAGCTTGCATGCCTCAGTCGACTTATTTATAAAGTTCATCCATGC-3’) to attach overhang- sequences for Gibson assembly. The destination vector pJB-CAT-ProD-mCherry (kindly provided by Robert Heinzen and Paul Beare, Rocky Mountain Laboratories, MT, USA) was digested with PstI and SalI. The Gibson assembly method was utilized to ligate the purified PCR product with the digested vector to create the *C. burnetii* expression vector pJB-CAT- ProD-tdTomato, which was subsequently used to electroporate and transform *C. burnetii* in ACCM-D medium as described before (Omsland et al., 2009; Schafer et al., 2017). Transformed cells were plated on ACCM-D agar plates containing 3µg/mL chloramphenicol (CA) and were incubated for 10 to 14 days. The screening for positive clones was done via PCR using primers a535 (5’-CACAGCTAACACCACGTCGTCC-3’) and a537 (5’-CTGCATCACTGGCCCATCGG-3’). Positive clones were expanded in liquid ACCM-D in presence of 3µg/mL CA and analyzed for fluorophore expression via fluorescence microscopy.

### Treatment of C. burnetii with itaconate

Five mL of ACCM-2 media were inoculated with *C. burnetii* NMII at a concentration of 10^6^ *Coxiella*/mL and propagated at 37°C, 5% CO_2_ and 2,5% O_2_. At different time points post- inoculation the bacteria were treated with ITA for the indicated time periods. As controls, PBS was added to the bacterial culture. For determination of colony forming units (CFU), the bacterial culture was centrifuged at 4,500 x g for 15 min and the pellet was resuspended in 200 µL ACCM-D. Ten µL of serial dilutions were dropped on ACCM-D/0,3% agarose plate in triplicates. The plates were incubated for 2 weeks at 37°C, 5% CO_2_ and 2,5% O_2_. CFU were calculated according to the corresponding dilution factor. To measure bacterial growth by optical density, *C. burnetii* cultures were propagated as mentioned above. At the indicated time points, 100 µL were directly taken from the culture to determine the optical density at OD_600_.

### Mouse macrophages

Primary bone marrow-derived macrophages (BMM) were propagated from the bone marrow of femurs and tibiae by culture in petri dishes in complete Dulbecco’s modified Eagle medium (cDMEM; DMEM [Life Technologies] plus 10% fetal bovine serum [FBS; Biochrom, Berlin, Germany], 50 μM β-mercaptoethanol, penicillin and streptomycin) complemented with 10% L929 cell-conditioned medium [LCCM] as a source of macrophage colony-stimulating factor (M-CSF) at 37°C, 5% CO_2_ and 21% O_2_ for 7 d. Non-adherent bone marrow cells were used after over-night culture in cDMEM with 10% LCCM, counted and plated in petri dishes at a density of 5 to 8 x10^6^ per 10-cm dish. On day 3 of culture, additional 5 ml of cDMEM with 10 % LCCM were added. Adherent macrophages were harvested on day 7 by treatment with Accutase (Sigma, Deisenhofen, Germany), washed and counted.

### Human monocyte-derived macrophages

Leukocyte Reduction System chambers were obtained from the Department of Transfusion Medicine at the University Hospital Erlangen (Ethics commission protocol number 111_12B). Peripheral blood mononuclear cells (PBMC) were separated using Biocoll separation solution and centrifugation for 30 min at 510 x g without brakes at room temperature. The interface containing PBMC was carefully removed, transferred to a fresh tube and washed once with PBS and centrifugation at 400 x g for 5 min at room temperature. After resuspending the pellet in PBS, platelets were removed by another centrifugation step at 90 x g for 10 min at room temperature. Next, CD14^+^ monocytes were isolated through positive selection using Miltenyi CD14 beads and LS MACS columns. Human monocyte-derived macrophages (MDM) were differentiated from purified monocytes by culture in complete RPMI 1640 containing 10% FBS, HEPES, penicillin/streptomycin, 50 µM mercaptoethanol, and 50 U/mL recombinant human GM-CSF (Leukine, Genzyme). On day 3 or 4 of culture, cells were fed with fresh media and GM-CSF, and then harvested on day 7 by incubation with Accutase diluted in PBS 1:4 ratio. After washing and counting, MDM were seeded in 96-well plates at a density of 1.5 or 2.0 x 10^5^ cells per well, and allowed to adhere and rest over-night in complete RPMI without antibiotics.

### In vitro infection of macrophages

The day before infection, murine BMM and human MDM were harvested, seeded and cultured at 37°C, 5% CO2 in DMEM (for BMM) or RPMI (for MDM) complete medium containing 10% fetal calf serum (FCS) and 50 µM β-mercaptoethanol without antibiotics. For microscopic analysis, 0.5 x 10^6^ macrophages were seeded on 10 mm coverslips in 24-well dishes, whereas in 96-well dishes a density of 1.5 x 10^5^ cells/well were used for qPCR of *C. burnetii* GE and mRNA expression of host genes. The cells were infected with NMII at a MOI of 10, if not indicated otherwise. Following infection, cells were incubated at 37°C and 5% CO2. After 4 h, cells were washed with cDMEM/cRPMI media without antibiotic to remove extracellular NMII and finally supplied with antibiotic free cDMEM/cRPMI. At the indicated time points, supernatants were removed and the cells lysed in PeqDirectLysis Buffer with Proteinase K (PeqLab, Germany) for DNA or TriReagent (Sigma, Deisenhofen, Germany) for RNA preparation, respectively.

### Immunofluorescence microscopy

For indirect immunofluorescence microscopy analyses, BMM were cultured on 10mm coverslips in 24-well dishes. At indicated points of time, cells were washed three times with equilibrated PBS, fixed for 15 min with equilibrated 4% paraformaldehyde (PFA) and permeabilized with ice-cold methanol for 1 min. Cells were quenched and blocked with 50 mM NH_4_Cl in PBS/ 5% goat serum (GS) for 60 min at room temperature. Incubation with primary antibody dilution in PBS/ 5% GS was conducted at room temperature for 60 min. Subsequently, cells were washed three times with PBS and further incubated with secondary antibodies in PBS/ 5% GS for 30 min at room temperature. After final 3 x washing with PBS, coverslips were mounted using ProLong Diamond containing DAPI. For visualization, a Carl Zeiss LSM 700 Laser Scan Confocal Microscope and the ZEN2009 software (Jena, Germany) were used.

In this study, we used primary antibodies directed against *C. burnetii* (polyclonal rabbit serum, Davids Biotechnologie, Regensburg, Germany) and LAMP-1 (monoclonal rat IgG, Developmental Studies Hybridoma Bank, Iowa, IA, USA). Secondary antibodies were labeled with Alexa Fluor 594 and Alexa Fluor 488, respectively (Dianova, Hamburg, Germany).

### In vivo infection experiments

All mouse experiments were approved by the regional government (Regierung von Unterfranken, animal protocols 54-2532.1-44/13 and 55.2.2-2532.2-854-14). Mice were bred at the PETZ of the University Hospital Erlangen and transferred to a biosafety level 2 animal room at least one week before infection. Sex-matched groups of mice were infected with NMII either intraperitoneally (5x10^7^ CFU/200µL PBS/mouse) or intratracheally (10^6^ CFU/50µL PBS/mouse), as described before (Kohl et al., 2019). Intratracheal infection was performed by direct injection of bacterial suspension into the trachea of the mice. To do this, mice were anaesthetized with isofluorane, placed in dorsal position and the trachea was accessed after incision of the skin. Prior to surgery, buprenorphine (0.1 mg/kg) was injected intraperitoneally for analgesia and mice were supplied with Butorphanol (0.12 mg/mL drinking water for 24h after the surgery). The physical condition of the mice was monitored regularly. At the indicated time points, mice were killed by cervical dislocation and organ tissue (∼20-50mg) of spleen, liver, and lung was collected. Tissue samples were preserved in PeqDirectLysis buffer (Peqlab, Erlangen, Germany) plus proteinase K (Roche) for DNA preparation or in RNAlater (Qiagen, Hilden, Germany) for subsequent RNA isolation. For histopathology, organ pieces were fixed in PFA 4% over night.

### DNA isolation and quantification of C. burnetii burden via qPCR

Organ tissue or macrophages in cell culture were lysed in PeqDirectLysis buffer plus proteinase K at 56°C under shaking overnight. Then, proteinase K was inactivated at 85°C for 45 minutes. DNA from organ tissue or macrophages was used directly for qPCR. We defined the ratio of *C. burnetii* genomic copies to BMM genomic copies as bacterial load per cell. To amplify *C. burnetii* DNA, a TaqMan-based quantitative PCR for the insertion sequence IS1111 was performed using 5’-CATCGTTCCCGGCAGTT-3’ as forward, 5’- TAATCCCCAACAACACCTCCTTA-3’ as reverse primer, and the internal fluorogenic probe 6FAM-CGAATGTTGTCGAGGGACCAACCCAATAAA-BBQ (TibMolbiol, Berlin, Germany). Host cell genomic copies were quantified from the same sample using a primer set specific for murine albumin gene (exon 7): forward, 5′-GGCAACAGACCTGACCAAAG-3′ and reverse, 5′-CAGCAACCAAGGAGAGCTTG-3′. For samples derived from human macrophages, genomic copies were determined using primers for the human albumin gene (forward, 5’-GTGAACAGGCGACCATGCT-3’, reverse 5’-GCATGGAAGGTGAATGTTTCAG-3’). Standard curves for human and murine genomes were generated by serial dilution of DNA from human PBMC or mouse spleen cells. To quantitate *C. burnetii* genome equivalents, a standard curve using titrated DNA prepared from a defined number of *in vitro* cultured NMII was included in each experiment. *C. burnetii* burden was calculated per cell by normalizing bacterial genome copy numbers to albumin copy numbers.

qPCR was carried out in 384-well optical plates on an ABI Prism 7900HT sequence-detection system. To quantify *C. burnetii* DNA, 2x Roche FastStart Universal Master Mix (6µL) with 0.2 µM final concentration of each primers and an internal fluorogenic probe 20 µM was used. For the quantification of host albumin gene we used 2x SYBRselect master mix together with 0.83 µM of each primer. For both qPCRs, 2 µL of isolated DNA (prediluted 1:4 for macrophage lysates and 1:20 for organ samples in ultrapure H_2_O) was used as template in a final volume of 12 μL per reaction.

### RNA isolation and gene expression analysis by qRT-PCR

RNA from organ tissue (stored in RNAlater stabilizing reagent until further processing) was isolated using peqGold TriFastTM (Peqlab) or TriReagent (Sigma-Aldrich, Deisenhofen, Germany) and cDNA was synthesized using High Capacity cDNA Reverse Transcription Kit (Applied Biosystems). Primers and probes were chosen from the Universal Probe library (Roche). The fold change in gene expression was determined by the ΔΔCT method, using HPRT as housekeeping gene for calculation of ΔCT values and samples of naïve wild type or heterozygous mice as calibrators. For human samples, PPIA was used as housekeeping gene and samples were calibrated to non-infected controls.

### Processing of organ tissue for histopathological analysis

After fixation of organ pieces for 24h in PFA 4% at 4°C tissue was washed with PBS and subsequently embedded in paraffin. 5 µm paraffin-embedded tissue slides were transferred to Star Frost microscope slides (Knittel GmbH, Braunschweig, Germany) for staining of the tissue.

### Immunohistochemistry staining of organ tissue

*C. burnetii* was detected using the α-*C. burnetii* antiserum described above. Detection of bound antibodies was performed with biotinylated secondary goat anti-rabbit antibody and ABC-Kit using the peroxidase substrate method DABImmPact (all from Vector Laboratories, Burlingame, CA, USA).

### Slide scanning and quantification of images

DAB-stained tissue arrays of *C. burnetii* infected mice were digitized using a Hamamatsu digital slide scanner, model C13210, with 40x magnification, automatic exposure and focussing, resulting in a resolution of 221 nm/pixel (114932 DPI). Tissues of interest were exported at 20x magnification as TIF-formatted tile images. Using the Colour Threshold method in Image J, the area under tissue and DAB-stained areas were recognized in the HSB colour space; as tissue: Hue 0°-360°, Saturation 4%-100% =, Brightness 64.5%- 100%; DAB: Hue 280°-70°, Saturation 25%-100%, Brightness 8%-100%. Binary masks were created. For smoothing, the masks were enlarged three times and then reduced again three times, each time by 1 pixel per step. Outliers with a radius of 5 pixels were removed. The positively coloured area of C*. burnetti* was calculated as the percentage of the DAB mask to the total tissue mask.

### GC-MS analysis

For GC-MS analysis, BMM were seeded (1 x 10^6^ cells / well) in 6 well-plates the day before infection in cDMEM without antibiotics. After overnight incubation, NMII was added at MOI-10. 4h after infection extracellular bacteria were washed away with cDMEM media, then supplied with ITA and its derivatives (4-OI and DMI). 24 hours after infection, BMM were washed 4 times with PBS (1ml/well), with careful removal of PBS to ensure complete removal of extracellular ITA and derivatives. Macrophages were lysed with ice-cold 80% methanol and lysates were immediately frozen at -80°C until further analysis. Sample preparation and measurement was performed as recently described (Hayek et al., 2019). The internal standard solution was expanded to include ^13^C_5_-ITA (Toronto Research Chemicals/Biozol, Eching, Germany) to ensure accurate ITA quantification. Total protein amount was determined using the fluorescent dye SERVA Purple (Serva, Heidelberg, Germany) as recently described (Berger et al., 2021).

### Flow cytometry analysis

BMM infected with fluorescent tdTomato-expressing *C. burnetii* were analysed by flow cytometry at the indicated time points. After discarding the media, cells were washed with PBS/2% FCS in 24- or 96-well plates according to the experimental setup. Then cells were fixed with 2% PFA for 30 minutes at room temperature. Following one more washing step, cells were gently scraped off the bottom of the plate in PBS/2% FBS with a pipette tip. Finally, cells were filtered through a 70 µm cell strainer into FACS tubes and analyzed using a BD LSRFortessa (BD Biosciences, San Jose, CA, USA). Single cells were gated and tdTomato fluorescence was quantified in the PE channel using FACS Diva and Flowjo software.

### MTT assay

MTT assay was performed to investigate cellular toxicity of ITA and its derivatives. At the indicated times, 20 µL of MTT solution (5 mg/mL) were added to each well of a 96 well-plate on top of the cells containing 100 µL medium. 3 to 5h after incubation at 37°C, 5% CO_2,_ 10% SDS in 0.01 N HCl (100 µL/well) was added to the MTT mixture and incubated overnight. Plates were measured by means of an ELISA reader at 550 nm; mean absorbance values of the wells containing media only (without cells) incubated with MTT and SDS solution were used as blank for calibration.

### Statistical analysis

Statistical analysis was performed as described individually for each graph in the figure legends using GraphPad Prism (version 8). P values <0.05 were deemed statistically significant and are indicated by an asterisk (*).

## Acknowledgements

We gratefully acknowledge Alexandra Schwarz and Dr. Roland Jurgons (PETZ) as well as Manfred Kirsch (Institute of Clinical Microbiology, Immunology and Hygiene) for animal care.

This work was done as part of the doctoral thesis of Md. Nur A Alam Siddique.

Funded by grants of the Deutsche Forschungsgemeinschaft (Collaborative Research Center 1181, project A6 to RL and AL and project C4 to US). JJ and KD were supported by the Bavarian Ministry of Science and the Arts within the Bavarian Research Network bayresq.net.

## Author contributions

Conceptualization: LK, NAS, AL, RL

Investigation: LK, NAS, BB, RB, AP, CD, MÖ, KTY, IH, MS, SR

Formal Analysis: LK, NAS, BB, SR, RL Methodology: MM, AL

Resources: JSL, US, AB, GK, PJM, SW, MY, VS, JJ, PO, DD, KP

Supervision: KD, AL, RL Writing – original draft: RL

Writing – review and editing: NAS, SR, PO, US, CB, AL, RL

## Conflict of interest

The authors declare no competing financial interests.

## The Paper Explained

### Problem

Chronic Q fever caused by the intracellular bacterium *Coxiella burnetii* is a severe infection with often lethal vascular complications. Antibiotics are frequently inefficient in chronic Q fever despite extended treatment courses. *C. burnetii* infects macrophages and replicates in the phagolysosome. The determinants of success or failure in the host immune response to *C. burnetii* are not well understood.

### Results

In this manuscript, we identify a host-protective role for the metabolite itaconate produced by the enzyme ACOD1 (aka IRG1) in macrophages in response to *C. burnetii*. ACOD1 mRNA was highly expressed in the spleens of mice after infection with *C. burnetii*, as well as in human and murine macrophages. Infected ACOD1-deficient mice developed high bacterial burden and significant weight loss. Treatment with itaconate corrected this phenotype in mouse and human macrophages, and protected mice after infection *in vivo*. Mechanistically, we show a strong direct inhibitory activity of itaconate on *C. burnetii*.

### Impact

These results establish ACOD1-derived itaconate as an essential macrophage effector molecule for containment of *C. burnetii* and resolving infection. Given the lack of efficient antibiotic regimens for chronic Q fever patients, our findings suggest itaconate as novel host-derived treatment strategy.

## Expanded View Figure legends

**Expanded View Figure EV1:** Immunofluorescence microscopy of *Acod1^-/-^* BMM 120 hours after infection with NMII at MOI 10. IFNγ was added or not 4 hours after infection when extracellular *C. burnetii* was removed. Staining for Lamp1 appears in green, staining for *C. burnetii* in pink, DAPI stain for DNA shows nuclei (examples marked by arrows) and large CCV in *Acod1^-/-^*BMM in the absence of IFNγ (marked by asterisk).

**Expanded View Figure EV2:** MTT assay performed 24 and 96 hours after addition of ITA, DMI and 4-OI to BMM infected with NMII. Each dot represents one mouse and data were pooled together from 2 individual experiments with ITA (n=4 mice per genotype) and one experiment with 250 µM 4-OI and 250 µM DMI (n=2 mice per genotype).

## List of abbreviations

(BMM) Bone marrow-derived macrophages

(ACOD1) Cis-aconitate decarboxylase 1

(NMII) Coxiella burnetii Nine Mile phase II

(CCV) Coxiella-containing vacuole

(DMI) Dimethyl-itaconate

(IRG1) Immune responsive gene 1

(ITA) Itaconate

(MDM) Monocyte-derived macrophages

(4-OI) 4-Octyl-itaconate

(TLR) Toll-like receptor

(SDH) Succinate dehydrogenase

## Notes

### Competing Interest Statement

The authors have declared no competing interest.

