## Supplemental figures for "Macrophages inhibit *Coxiella burnetii* by the ACOD1-itaconate pathway for containment of Q fever"

**Kohl et al. Expanded View Figures**

### Expanded View Figure EV1:
### Immunofluorescence microscopy of *Acod1*$^{-/-}$ BMM treated with IFNγ.

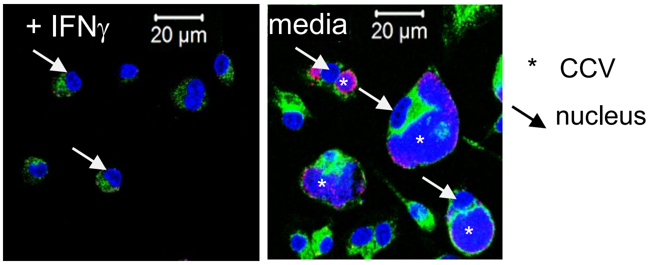


**Expanded View Figure EV1:** Immunofluorescence microscopy of *Acod1*$^{-/-}$ BMM 120 hours after infection with NMII at MOI 10. IFNγ was added or not 4 hours after infection when extracellular *C. burnetii* was removed. Staining for Lamp1 appears in green, staining for *C. burnetii* in pink, DAPI stain for DNA shows nuclei (examples marked by arrows) and large CCV in *Acod1*$^{-/-}$ BMM in the absence of IFNγ (marked by asterisk).

Kohl et al. Expanded View Figures

**Expanded View Figure EV2: MTT Assay of BMM treated with ITA, 4-OI and DMI.**

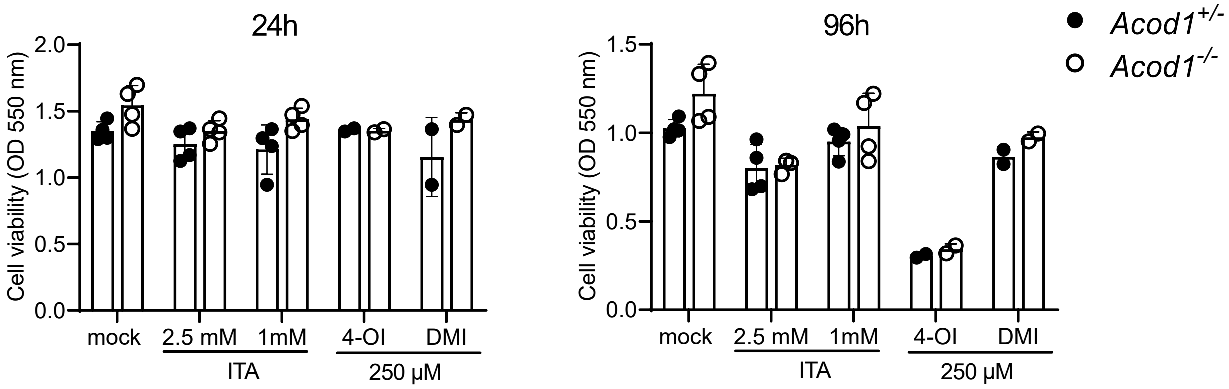


**Expanded View Figure EV2:** MTT assay performed 24 and 96 hours after addition of ITA, DMI and 4-OI to BMM infected with NMII. Each dot represents one mouse and data were pooled together from 2 individual experiments with ITA (n=4 mice per genotype) and one experiment with 250 μM 4-OI and 250 μM DMI (n=2 mice per genotype).
